# A comprehensive review of South Australia’s Great Artesian Basin spring and discharge wetlands biota

**DOI:** 10.1101/2023.10.29.564639

**Authors:** P. G. Beasley-Hall, B. A. Hedges, S. J. B. Cooper, A. D. Austin, M. T. Guzik

## Abstract

**Context:** The Great Artesian Basin (GAB) feeds thousands of springs in Australia’s arid centre, supporting relictual species not found elsewhere on Earth. Springs are considerably threatened by ongoing water abstraction by industry. Robust management plans are needed to prevent further extirpations of GAB taxa, but fundamental biodiversity knowledge is lacking.

**Aims:** We aimed to characterise major organismal groups in South Australian GAB springs and surrounding wetlands, their conservation and taxonomic status, and potential biodiversity hotspots and connectivity of spring ecosystems.

**Methods:** Focusing on South Australia as a case study, we conducted a comprehensive review of GAB spring biota based on the published scientific and grey literature.

**Key results:** Almost 500 taxa have been recorded from GAB springs, the majority being invertebrates. Community composition is highly heterogeneous among spring clusters and the true extent of spring biodiversity is far greater than currently known.

**Conclusions:** GAB springs have intrinsic value as refugia for both endemics and cosmopolitan taxa. GAB invertebrates are poorly conserved and largely lacking in taxonomic knowledge. We highlight several potential biodiversity hotspots that have been overlooked in the literature.

**Implications:** Fundamental biodiversity information on the GAB is crucial for decision-making in conservation management, for industry, and for Traditional Custodians.

**Lay summary:** The Great Artesian Basin (GAB) is Australia’s largest freshwater resource. Springs fed by the GAB support many species not found elsewhere on Earth, but conservation is hindered by a lack of fundamental knowledge about the plants, animals, and fungi reliant on these habitats. Using South Australia as a case study, we provide a comprehensive review of GAB biodiversity in that state.

## 1 Introduction

The Great Artesian Basin (GAB) is Australia’s largest groundwater resource, spanning over one fifth of the continent’s area or almost 2 million km^2^ (Habermehl, 2020). At the outer margins of the GAB where the confining layer is thin, pressurised water is often forced to the surface to form springs and associated wetlands. Thousands of such springs are found throughout Central Australia, with >5,000 individual spring vents (discrete discharge points of water) in South Australia (hereafter SA) (Arabana Aboriginal Corporation, 2021; Government of South Australia, 2023), >2,000 in Queensland (Queensland Department of Regional Development, Manufacturing and Water, 2023), and >400 in New South Wales (New South Wales Department of Planning, Industry and Environment, 2021). From an ecological and evolutionary standpoint, GAB-fed springs support wetlands representing “museums of biodiversity” housing plant and animal species not found elsewhere on Earth (Murphy et al., 2015). As relicts of the continent’s mesic past, species endemic to GAB springs often have exceptionally small distributions. It is not uncommon for taxa to be restricted to a single cluster of springs, termed *ultra-short range endemics* (examples found in Gotch et al., 2008; Guzik et al., 2019, 2012; Harvey, 1990; King, 2009; King et al., 2014; Murphy et al., 2012, 2009).

The conservation and management of groundwater resources is lacking globally (Famiglietti, 2014). Springs fed by the GAB are no exception and are considerably threatened by a range of industrial and pastoral practices (Fairfax and Fensham, 2002; Lewis and Harris, 2020; Mudd, 2000). The sinking of more than 50,000 artificial boreholes and the direct abstraction of Basin water have led to substantial reductions in artesian pressure (hereafter *drawdown*) and spring flow (Beasley-Hall et al., 2023c; Gotch et al., 2016; Great Artesian Basin Coordinating Committee, 2019; Mudd, 2000). A complete cessation of flow has occurred for an estimated 800 springs across Australia (Andersen et al., 2016; Fensham et al., 2016), and declines of endemic fauna have been documented as a result of this degradation of spring habitats (Fensham et al., 2010). Springs and their biota also face threats in addition to drawdown, including grazing, trampling, and soiling of wetlands by livestock such as cattle (Fatchen, 2000), overabundant pest species (Kerezsy, 2015; Noack, 1994), climate change (Ordens et al., 2020), and tourist activity (Witjira National Park Co-management Board, 2022). Despite remediation efforts, drivers of spring flow remain poorly understood and net water flows in GAB springs appear to still be reducing in certain regions of the Basin (Green and Berens, 2013).

The development of a biotic inventory has been identified as an urgent action to recover the community of native species dependent on GAB springs (Fensham et al., 2010). Publicly available data can help to inform future management plans and early warning systems that detect changes in these ecosystems of high biodiversity value (Brack et al., 2015; Obura et al., 2019; Vaughan et al., 2001) and so improving their accessibility is a clear priority. Further, industry stakeholders frequently rely on biotic inventories to ensure they are fulfilling their environmental obligations. This is particularly relevant for the GAB in the context of drawdown. Yet, basic taxonomic and biological information for these communities is still lacking, inaccessible, or disparate, making it difficult to accurately assess and monitor all spring species in the face of rapid, human-driven change. The Queensland Government has published information on metrics such as water quality, chemistry, flow rate, biodiversity, and spring condition (Queensland Government, 2018), but such information is not public for SA and New South Wales. This lack of taxonomic information is particularly pronounced for endemic spring invertebrates (Rossini, 2020). The relationship between environmental characteristics of springs and biodiversity is also not well understood (Fensham et al., 2010; Green and Berens, 2013; Harris, 1992; Rossini et al., 2018), meaning it is currently difficult to establish clear conservation priorities for GAB spring taxa in certain locations over others. The compilation of baseline information on these taxa in a centralised, accessible manner would represent a fundamental resource to facilitate future conservation work on this ecological community. Here, we conducted a literature review of species associated with springs in South Australia, selected due to its high number of GAB springs (>60% of all active springs) and the comparatively well-studied nature of its biodiversity (King, 2009; Murphy et al., 2015, 2013; Zeidler and Ponder, 1989). We specifically aimed to: 1) Characterise major organismal groups in SA GAB springs and surrounding wetlands; 2) Document the conservation and taxonomic status of these groups to highlight knowledge gaps; 3) Identify hotspots of biodiversity in SA GAB springs; and 4) Assess connectivity of SA GAB spring communities.

## 2 Materials and Methods

### 2.1 Literature review and database construction

We undertook a non-systematic review of all available information relevant to the biodiversity of GAB springs in South Australia. A formal systematic searching strategy was not possible as a large proportion of information on springs is present in unpublished or grey literature such as government reports, internal publications from mining companies, and museum records. We therefore relied heavily on repositories such as the Government of South Australia’s WaterConnect portal (South Australian Department for Environment and Water, 2023) and The Atlas of Living Australia (Belbin et al., 2021). We were specifically interested in data indicating the presence or absence of species in or around spring wetlands as well as their occurrence extents, if available, to gauge where taxa occurred and their degree of endemicity. The scope of this review spanned the three spring supergroups in SA: Kati Thanda–Lake Eyre, Munda / Lake Frome, and Dalhousie (Figure 1). **Supergroups** are clusters of **spring complexes**, which themselves contain **spring groups** composed of individual **springs**. Spring complexes share similar geomorphological characteristics and water chemistry, whereas groups are clusters of springs sourcing from the same fault or structure (Lewis et al., 2013). Individual springs are composed of permanent wetland vegetation with at least one **vent**, a discrete discharge point of water (Gotch, 2013). Within these three supergroups, we focused on 23 spring complexes containing 170 spring groups (Beasley-Hall et al., 2023a for raw research data via FigShare). Whilst the number of these spring clusters, as well as their naming conventions, have previously not been standardised (Gotch et al., 2016), we selected these locations due to their widespread use in conservation management by the Australian Government (Department of the Environment, 2022; Lewis and White, 2013).

**Figure 1:**
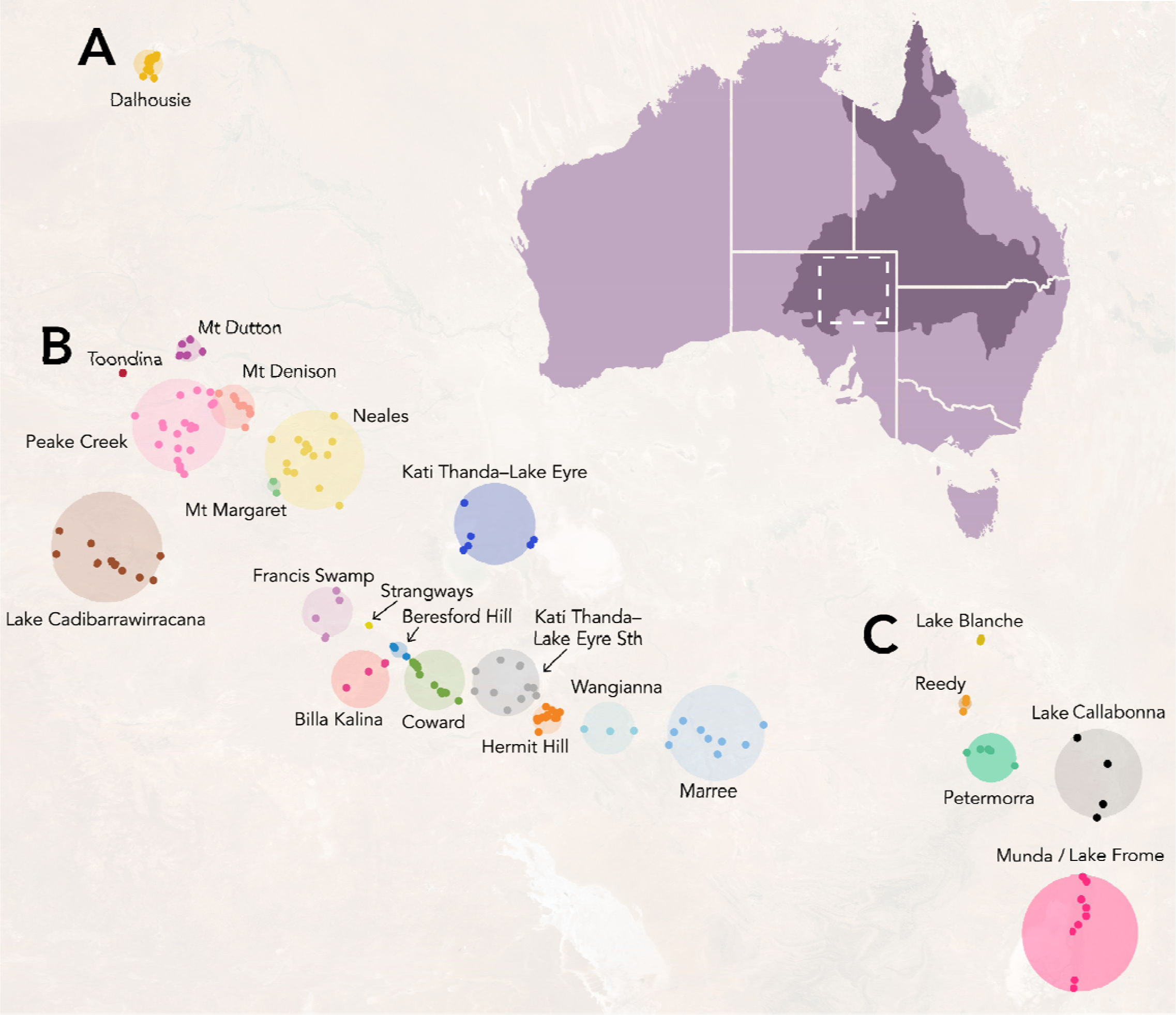
Springs fed by the Great Artesian Basin (GAB) in South Australia. Springs are classified hierarchically as supergroups, the broadest classification (A, Dalhousie; B, Kati Thanda–Lake Eyre; C, Munda / Lake Frome); complexes (labelled coloured circles); groups (coloured points); springs; and finally vents, discrete water discharge points. The approximate area of the GAB is shown in dark purple and the location of the SA GAB springs is indicated by the dashed square.

We retrieved Aboriginal names for spring groups from a key anthropological study (Hercus & Sutton, 1985) to ensure locations were referred to by a dual naming convention whenever possible. The validity of these names was confirmed by The Arabana Aboriginal Corporation in a personal communication to the authors. For locations with dual names, but lacking “official” formatting (e.g., not used by federal/state government legislation or publications), we followed the Australian Government’s style manual (Australian Government, 2023). We chose not to standardise names across springs, groups, or complexes as they refer to specific locations with distinct associated mythologies and histories (Hercus & Sutton, 1985). Information below the spring group level, where present, was standardised to the group level due to the rarity of such records and the inconsistency of spring vent naming conventions. As GAB spring fauna are often morphologically cryptic yet genetically distinct, we recorded species as separate taxa if explicitly indicated by their species authorities based on genetic data following Rossini *et al*. (2018).

To supplement presence/absence records of spring taxa, we also gathered information related to taxonomic status, conservation status, common names, synonyms, and endemicity where available. Taxonomic status was coded into three categories: taxa for which there is a corresponding formal taxonomic description (“described”), taxa awaiting taxonomic description (“undescribed”), or those for which species-level occurrence records were not available (e.g., a family-only record; “unidentified”). For taxa with species-level identifications, conservation status information was retrieved from the Species Profile and Threats Database to capture listings under the IUCN Red List of Threatened Species and Australian federal and state environmental legislation (Department of Climate Change, Energy, the Environment and Water, 2023). Finally, for each endemic taxon we noted whether it occupied only one spring group, complex, or supergroup. Except for undescribed species with well-established occurrence records, endemicity was not recorded for taxa without a species-level identification.

### 2.2 Biodiversity metrics

For each spring group, we first transformed presence/absence records to weight them by whether they represented resolution at the spring group level (hereafter confident records) or those which referred to a taxon’s presence within a certain complex but did not supply spring group information (hereafter coarse records). To assess the inclusion of uncertain occurrence information, we calculated metrics for our entire dataset and a subset of the data only considering confident records. As the dataset included only presence/absence records, we were limited in our choice of biodiversity metrics and focused on species richness and endemicity. We calculated species richness values (hereafter taxon richness) by the number of putative taxa in each spring. Spring groups were ranked by the degree of endemicity of their biota using a modification of the scoring system developed by Fensham and Price (Fensham and Price, 2004). Originally applied to GAB flora in spring complexes in Queensland, the ranking has since been expanded to fauna across Australia (Rossini et al., 2018) and relies on the number of populations corresponding to the most widespread taxon in a spring dataset. The desert goby *Chlamydogobius eremius* (Zietz, 1896) is the most widespread SA GAB spring endemic, occurring across 24 known groups (Gotch et al., 2016; Rossini et al., 2018); we used this value as a proxy for the taxon’s number of populations, although we acknowledge this may be an underestimate as spring groups do not necessarily share permanent wetlands. Rankings for each endemic taxon were first calculated by dividing 24 by the number of groups the taxon occurred in such that *C. eremius* (24/24) would receive the lowest ranking due to its comparably widespread distribution. As all endemic taxa assessed here do not occur beyond their respective supergroups, each taxon was then scored by whether it was further restricted to a single spring complex (+1) or group (+2). These scores, hereafter *endemicity rankings*, were summed for each spring group. We visualised these metrics using QGIS (QGIS Association, 2023). Spring groups were first mapped using latitude and longitude information corresponding to vents retrieved from the Government of South Australia’s WaterConnect portal (Government of South Australia, 2023); for groups containing multiple vents, centroids were calculated to approximate their location. Circles corresponding to the above metrics per group were scaled using the Flannery method. To assess differences in community composition among spring groups, we calculated pairwise Jaccard distances using a binary matrix of presence/absence records using the *proxy* package in R v.4.3.0 (Meyer and Buchta, 2022; R Core Team, 2023). Principal components analysis was performed using the native R *stats* package and visualised using *ggplot2* (R Core Team, 2023; Wickham, 2016). Springs without occurrence records were excluded from the analysis, as were taxa known to occur in SA GAB springs generally but without specific location information. Finally, we produced rarefaction curves using the R package iNEXT (Hsieh and Chao, 2022) from our entire dataset and a subset of the data considering only confident records. Rarefaction was performed using the rarefaction and extrapolation models for species richness (q = 0), 95% confidence intervals, and 100 replications.

## 3 Results

### 3.1 Biodiversity of the SA GAB

The database we compiled based on our comprehensive literature review captured 3,463 occurrence records corresponding to 495 distinct and putatively distinct taxa. Of these records, 2,300 were considered coarse—i.e., corresponding to the supergroup or complex level, but not informing the presence of taxa at a specific spring group. Invertebrates were by far the most speciose group in the dataset (42%, 207 taxa) followed by vascular plants (22%, 111 taxa), vertebrates (21%, 102 taxa), algae (14%, 68 taxa; for the purposes of this paper, includes green, golden, and blue-green algae) non-vascular plants (1%, 5 taxa), and fungi (0.4%, 2 taxa) (Figure 2). Sixty-five taxa (13%) are known only from SA GAB springs, almost all of which are invertebrates. Despite the species diversity dominance of invertebrates, these organisms are also the most poorly known in GAB springs from a taxonomic standpoint. Just over one-third of invertebrate taxa have formal taxonomic names (37%), whereas the remainder of the fauna is either undescribed (14%) or has an unknown taxonomic status due to a lack of species-level occurrence records (49%). For other taxonomic groups, taxonomic description ranges from 50–100% (described), 0–1.5% (undescribed), and 0–50% (unidentified). Further, apart from the Gastropoda, no invertebrate taxon has had its conservation status assessed at the global, federal, or state level (Figure 2). Fifty-eight of the 170 spring groups assessed in this study had no corresponding occurrence records in the literature. A rarefaction curve derived from the dataset suggests the artesian wetlands of SA have not been adequately surveyed, and if additional locations were examined with equal sampling intensity, dozens of additional taxa would likely be documented (Figure 3).

**Figure 2:**
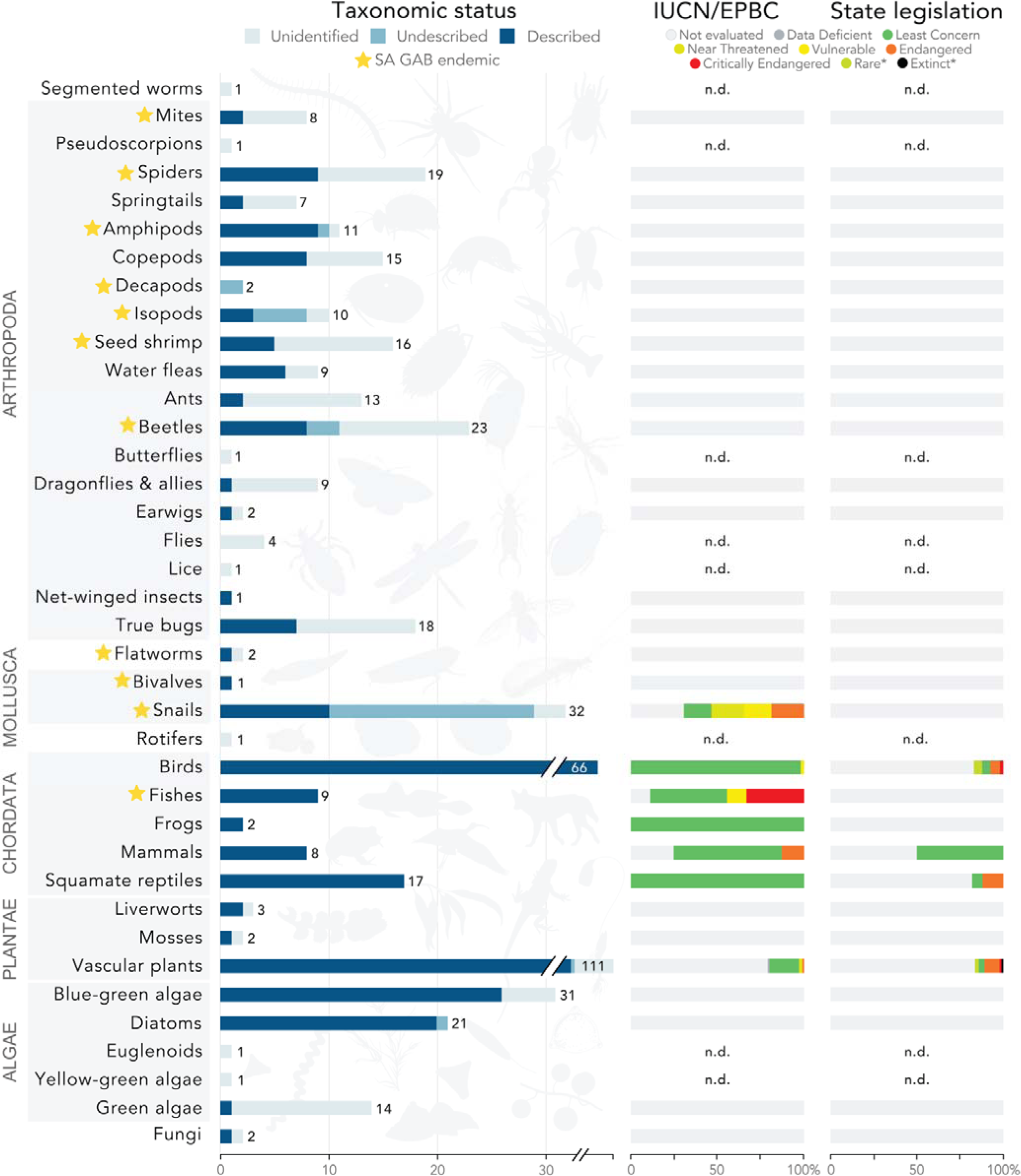
Fauna, flora, and funga associated with SA GAB spring wetlands, with major organismal groups indicated by grey boxes. Groups lacking species-level records have an unknown conservation status and are indicated with n.d. (no data). * Rare and Extinct are specific to SA/TAS and NSW legislation, respectively, and only refer to populations/taxa within those states. Silhouette credits Maxime Dahirel, Armelle Ansart, Mathieu Pélissié, and Lafage via PhyloPic.

**Figure 3:**
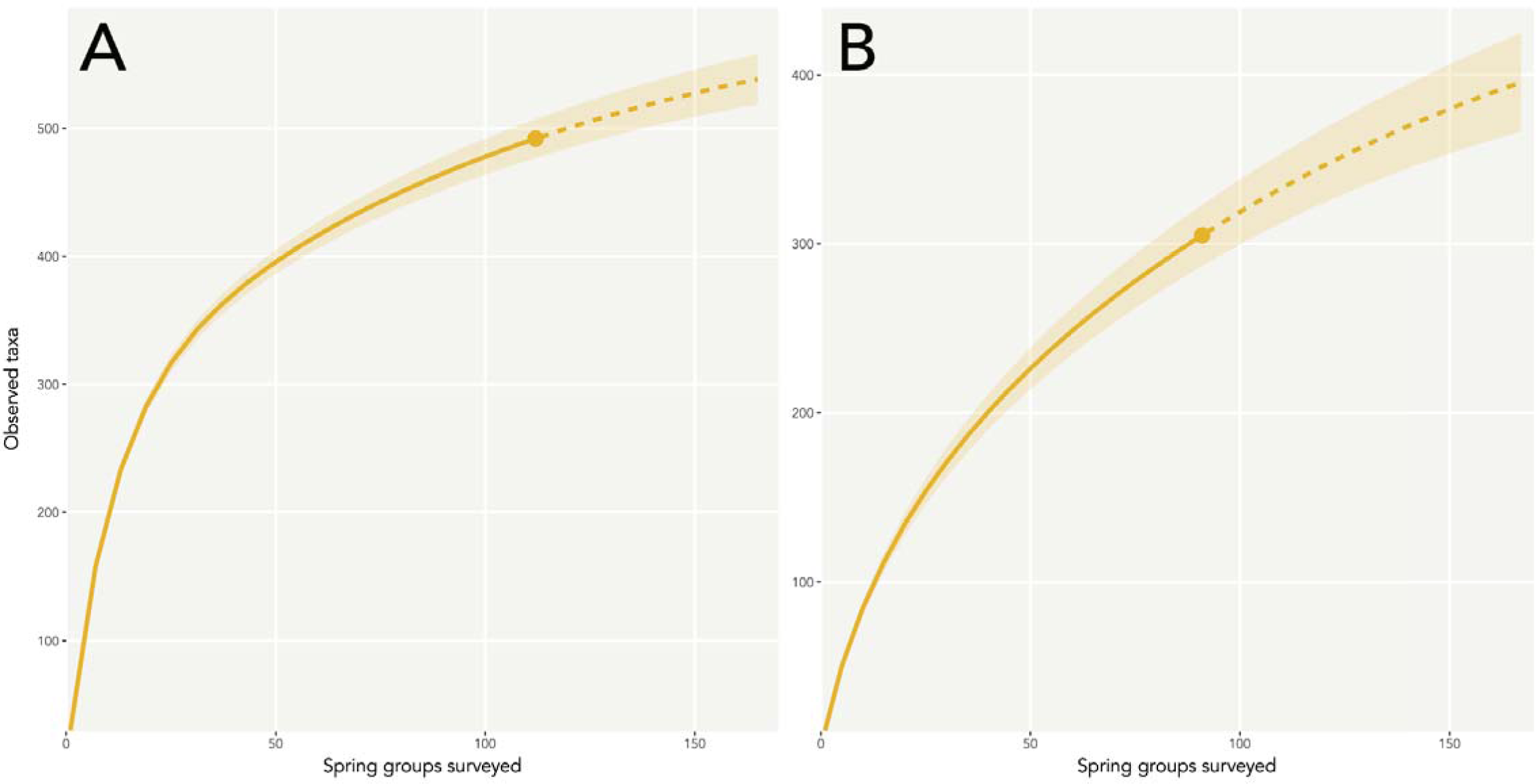
Extrapolation curves predicting an increase in taxon richness if additional spring groups were surveyed, calculated from all occurrence records in the dataset (A) and a subset of the dataset containing only confident records (B). As the curve does not flatten at the maximum number of known spring groups in the state (170), an increase in sampling effort *within* springs (as opposed to the sole sampling of additional springs) may also capture a more comprehensive representation of these communities. Dotted lines represent extrapolation beyond the taxon richness of our dataset (solid lines); shaded areas represent 95% confidence intervals.

In addition to occurrence records in the dataset, we also identified several records of potential local extinctions in the literature (Table 1). In all, we retrieved absence records corresponding to one isopod taxon, which may represent multiple species (*Phreatomerus latipes* (Chilton, 1922) Central, North, South haplotypes) (Fensham et al., 2010; Guzik et al., 2012; Kinhill, 1997; Kinhill-Stearns, 1984), one ostracod (*Ngarawa dirga* De Deckker, 1979) (Fensham et al., 2010; Kinhill, 1997; McLaren et al., 1985), 15 snail taxa (*Fonscochlea accepta*, members of *F. aquatica*, *F. billakalina*, *F. variabilis*, *Trochidrobia punicea*, *T. smithi* species complexes [all Ponder, Hershler & Jenkins, 1989], *Sinumelon pedasum* Iredale, 1937) (Department of the Environment, 2022; Fensham et al., 2010; Ponder et al., 1995; Rossini et al., 2018; Zeidler and Ponder, 1989), and four fishes (the Dalhousie goby *Chlamydogobius gloveri* Larson, 1995, Lake Eyre hardyhead *Craterocephalus eyresii* (Steindachner, 1883), spangled perch *Leiopotherapon unicolor* (Günther, 1859), Dalhousie gudgeon *Mogurnda thermophila* Allen & Jenkins, 1999) (Department of the Environment, 2022; Froese and Pauly, 2023; Gotch et al., 2016; Kodric-Brown et al., 2007; Rossini et al., 2018; Zeidler and Ponder, 1989). To the best of our knowledge, these records corresponded to likely local extinctions as opposed to complete species extinctions (i.e., relevant taxa were present in at least one additional location). These likely extirpations have occurred across 13 spring groups in the Dalhousie and Kati Thanda– Lake Eyre supergroups (Table 1). Given the patchy sampling of GAB springs during many surveys, we note that these data may be a result of stochastic sampling effort rather than true extirpations and is therefore an opportunity for future research. However, we note for certain taxa this evidence is more robust than others, e.g. the discovery of empty snail shells at springs (Ponder et al. 1989).

**Table 1:** Records of likely local extinctions of fauna in the SA GAB springs. To our knowledge there are no records of floral extinctions in the SA GAB. ^ Taxon represents multiple genetically distinct clades which we consider putative species, but which of these were present in the below location(s) prior to their apparent extirpation is unknown. KT–LE Sth = Kati Thanda–Lake Eyre South. * GAB spring endemic.

| **Taxon** | **Broad grouping** | **Spring group(s)** | **Spring complex(es)** |
| --- | --- | --- | --- |
| *Phreatomerus latipes*<sup>^*</sup> | Isopod | Venable, Marrinha / Hergott | Hermit Hill, Marree |
| *Ngarawa dirga*<sup>*</sup> | Ostracod | Venable, Manda-wardunha / Mundowdna | Hermit Hill, Marree |
| *Fonscochlea accepta*<sup>*</sup> | Hydrobiid snail | Venable, Palura Pintjanha / Priscilla | Hermit Hill, KT–LE Sth |
| *Fonscochlea aquatica*<sup>^*</sup> | Hydrobiid snail | Margaret | Francis Swamp |
| *Fonscochlea billakalina*<sup>*</sup> | Hydrobiid snail | Margaret | Francis Swamp |
| *Fonscochlea variabilis*<sup>^*</sup> | Hydrobiid snail | Venable, Palura Pintjanha / Priscilla | Hermit Hill, KT–LE Sth |
| *Fonscochlea zeidleri*<sup>^*</sup> | Hydrobiid snail | Venable, Palura Pintjanha / Priscilla, Centre Island | Hermit Hill, KT–LE Sth |
| *Trochidrobia punicea*<sup>^*</sup> | Hydrobiid snail | Venable, Palura Pintjanha / Priscilla | Hermit Hill, KT–LE Sth |
| *Trochidrobia smithi*<sup>^*</sup> | Hydrobiid snail | Margaret | Francis Swamp |
| *Sinumelon pedasum* | Camaenid snail | Irrwanjira / Errawanyera | Dalhousie |
| *Chlamydogobius gloveri* | Fish | Irrwanjira / Errawanyera, Frog Dreaming, Kirki /<br>Dalhousie Proper, Cadni Dreaming | Dalhousie |
| *Craterocephalus eyresii* | Fish | Old Nilpinna, Thuntinha / Toondina | Peake Creek, Toondina |
| *Leiopotherapon unicolor* | Fish | Idnjundura / Kingfisher, Ilpikwa | Dalhousie |
| *Mogurnda thermophila* | Fish | Frog Dreaming | Dalhousie |

### 3.2 Biodiversity metrics and community composition

Taxon richness was more heavily impacted by the exclusion of coarse occurrence records—as most data in this category are not spring endemics—whereas endemicity rankings were essentially unchanged between spring locations. Complexes may also be taxon-rich without containing a large number of endemics (e.g., groups within the Francis Swamp complex; Figure 4) and vice versa (e.g., Beresford Hill). Groups with high levels of richness values generally corresponded to the Coward, Dalhousie, Billa Kalina, and Hermit Hill complexes and those containing large numbers of isolated endemics included Dalhousie, Francis Swamp, Mount Denison, and Coward (Figure 4; (Beasley-Hall et al., 2023b) for raw research data via FigShare).

**Figure 4:**
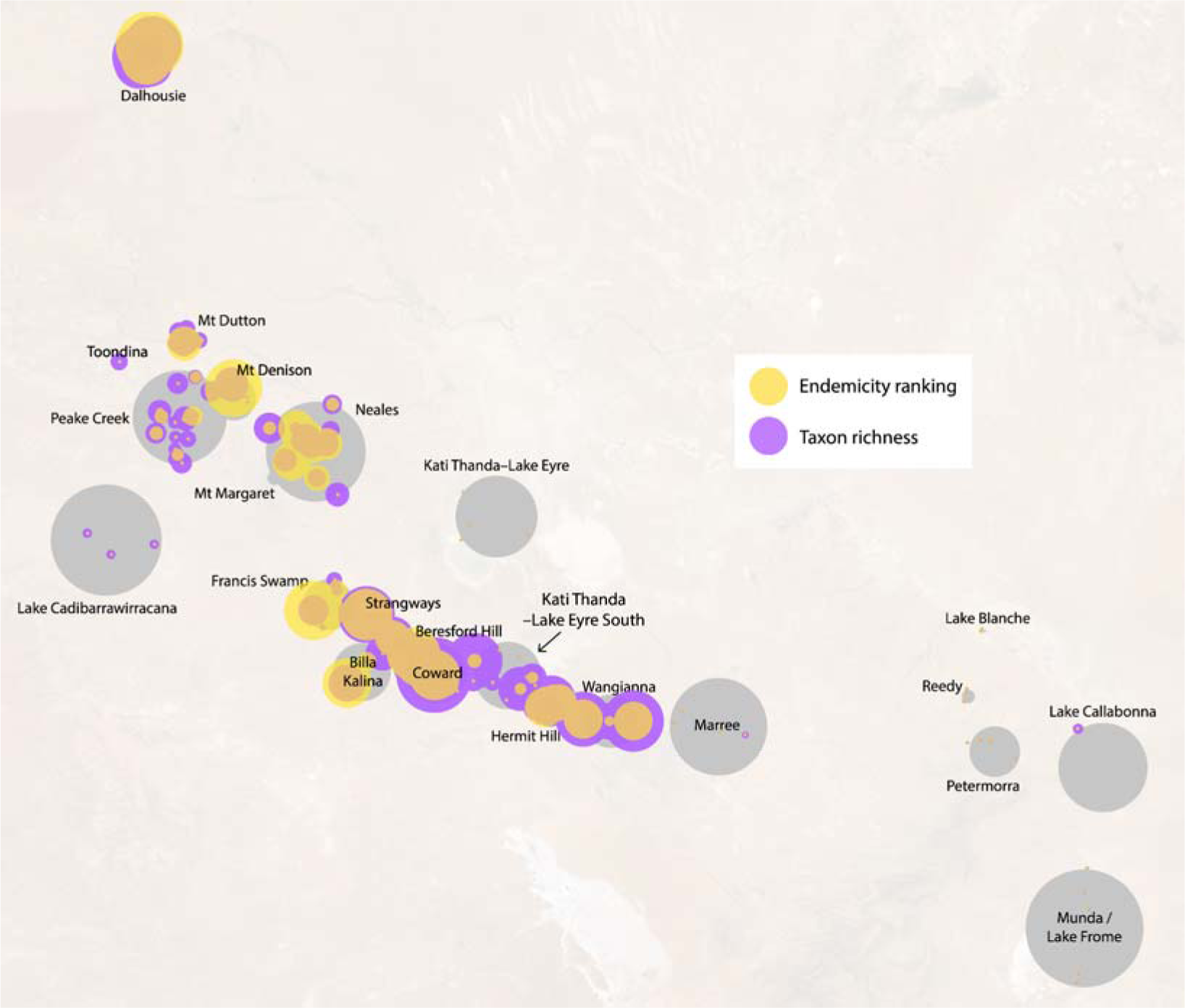
Rankings of endemicity and overall biodiversity in SA GAB spring groups, with circle size indicating a greater value for a given metric. Larger circles indicate a higher proportion of endemics and/or more isolated endemic fauna (endemicity ranking, yellow) or a greater number of putative species (taxon richness, purple). Grey circles represent the approximate area of spring groups as shown in Figure 1. This visualisation was calculated from both coarse and confident records; raw data, including metrics calculated from only “confident” records, are available in Supplementary File S2.

While there is some degree of overlap regarding taxon composition of spring complexes, particularly in the Kati Thanda–Lake Eyre region (Figure 5), locations with similar biodiversity metrics do not necessarily support the same biotic communities. The Peake Creek, Lake Cadibarrawirracanna, Neales River, and Mount Dutton complexes of the Kati Thanda–Lake Eyre supergroup are generally non-overlapping and groups from within the Dalhousie complex/supergroup are easily differentiated from other complexes (Figure 5). In contrast, groups from the Munda / Lake Frome complex are nested within those from the Kati Thanda–Lake Eyre supergroup. Groups belonging to the same complex are also generally more similar to one another than those in different complexes. This apparent lack of connectivity is perhaps unsurprising given the aridity of the surrounding landscape. Overall, spring groups within the Dalhousie supergroup/complex are the most distinct in the dataset in that they support taxa not known to be associated with other spring complexes, including highly isolated endemics (Figures 4, 5).

**Figure 5:**
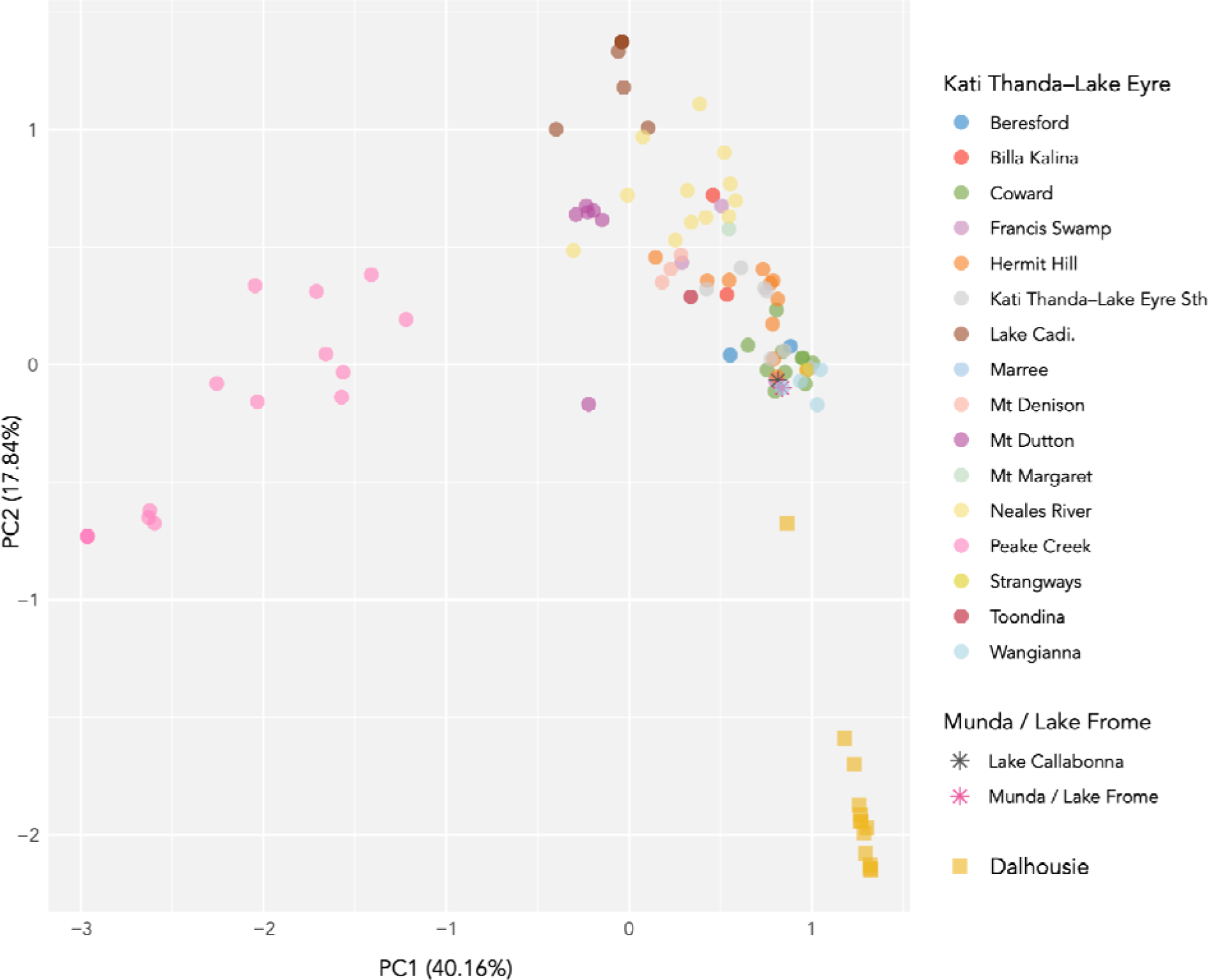
Principal components analysis (PCA) showing composition of faunal, floral, and fungal communities reliant on South Australian GAB springs. Both coarse and confident occurrence records in the dataset were used to calculate Jaccard distances between spring groups. Spring groups lacking occurrence records are not shown here. Groups are coded by their respective spring complex (colour) and supergroup (shape). Axis labels refer to the amount of variability explained by each principal component (PC).

## 4 Discussion

Here, we present a robust literature review of the biota associated with artesian wetlands of South Australia. These data highlight several key trends: 1) invertebrates are a poorly known component of GAB biodiversity relative to the rest of the biota; 2) the composition of GAB biota differs considerably by location, challenging previous conceptions of relevant biodiversity hotspots; and 3) the true extent of GAB biodiversity is clearly far greater than is currently appreciated in the literature. We discuss each of these findings below, as well as conservation implications for GAB springs more broadly and how a centralised biodiversity resource for the GAB could be implemented in future.

### 4.1 Biodiversity of South Australia’s GAB springs

Almost 500 taxa are associated with GAB springs and their surrounding wetlands. Invertebrates represent the largest proportion of this biota and the largest group of endemics, with the majority of being insects, crustaceans, and arachnids (Figure 2). Invertebrates are poorly documented from biodiversity, ecological, and conservation standpoints relative to the remainder of GAB biota and are also overrepresented in extinction records (Table 1). These groups may therefore be the most at risk of decline due to difficulties in devising management strategies. In contrast, while vertebrates were the third-most speciose group associated with the system (N = 102), almost all were widespread species not of conservation concern (Fensham and Price, 2004) (Figure 2). However, we note here that vertebrates such as waterbirds are nonetheless dependent on the springs as breeding grounds (Badman, 1985), and may make more extensive use of these habitats in non-flood years and/or as “stepping stone” habitats to sustain migrations to nearby Kati Thanda–Lake Eyre. For example, the little grassbird *Poodytes gramineus* (Gould, 1845) is not restricted to GAB springs, but nonetheless has an obligate relationship with stands of the reed *Phragmites australis* in GAB wetlands (Read, 1997).

Our data suggest a high degree of heterogeneity in the ecological communities of the SA GAB and challenge past assumptions regarding the system’s biodiversity hotspots. It has long been recognised that certain SA GAB springs are more speciose than others, but only recently has a quantitative view of candidate biodiversity hotspots been put forward (Rossini et al., 2018), and without consideration of non-endemic taxa. Springs at Dalhousie are one such hotspot proposed by Rossini *et al*. (2018), which we support in our review here. We also propose the Francis Swamp, Mount Denison, and Coward complexes as hotspots due to their contribution to the remaining bulk of endemicity in the SA GAB. Indeed, these springs house a much more speciose and isolated endemic biota than initially recognised based on published records (Supplementary File S1). Yet, we note that not all taxon-rich locations are necessarily rich in endemic species (Figure 4). Overall, our findings indicate the disturbance or extinction of any given spring group could represent an enormous loss of biodiversity when both endemic and non-endemic taxa are considered together. The biodiversity of the South Australian GAB has not been adequately surveyed (Figure 3) and over two-thirds of records we collated were coarse, meaning they lacked occurrence information at the spring group level. While records are helpful in distinguishing spring group locations and suggest considerable structure in taxa occupying different spring complexes and supergroups (Figure 5), they nonetheless highlight a lack of detailed past surveys and the sparseness of published information on GAB biodiversity.

The observed variability in the dataset may also be associated with sampling bias. Spring groups with the highest endemicity rankings, such as those within the Dalhousie and Coward complexes, are also those consistently associated with high sampling effort in previous surveys (Badman, 1985; Greenslade, 1985; Kovac, 2003; Mitchell, 1985; Noack, 1994; Ponder et al., 1995; Sokol, 1987; Zeidler and Ponder, 1989). The Kati Thanda–Lake Eyre and Dalhousie supergroups corresponded to over 2,000 occurrence records, whereas observations within the Munda / Lake Frome supergroup were restricted to a single, coarse occurrence record of the copepod *Microcyclops dengizicus* (Lepeschkin, 1900) (Zeidler and Ponder, 1989). Munda / Lake Frome has clearly not been adequately sampled, although this does not necessarily imply the location lacks endemic taxa yet to be characterised (McLaren et al., 1985; Rossini, 2020). Indeed, widespread taxonomic groups expected to be present in springs, such as the Hymenoptera and other cosmopolitan insect groups, were also poorly represented in the dataset due to a presumed bias in sampling methods and a lack of past taxonomic expertise. As such, we caution against the derivation of conservation priorities from locations with high measures of metrics such as species richness alone, but trends are nonetheless evident in this review regarding spring locations and taxonomic groups that have been *under*sampled.

### 4.2 Conservation implications for South Australia’s artesian springs

At least 14 taxa may have become locally extinct in the SA GAB springs (Table 1). Evidence for extirpations includes survey work conducted prior to the extinction of springs themselves or, conversely, the discovery of remains at extinct springs (e.g., snail shells). The bulk of these likely extirpations have occurred in spring groups that have ceased to flow because of drawdown, namely Venable, Palura Pintjanha / Priscilla, Marrinha / Hergott, Margaret, and Manda-wardunha / Mundowdna (Fensham et al., 2010). The Venable and Palura Pintjanha / Priscilla spring groups became extinct around 1990 following predictions by industry that mining activities would lead to a partial, if not complete, reduction in artesian flow at certain GAB springs (Kinhill-Stearns, 1983, 1982). Marrinha / Hergott ceased to flow in the mid-1980s as water was withdrawn to supply the nearby town of Marree (McLaren et al., 1985, p. 198). Other extirpations have been attributed to human modification of springs, the presence of overabundant invasive and native species, and potentially insufficient sampling effort. The apparent local extinction of the fish species, Lake Eyre hardyhead (*Craterocephalus eyresii*) at Toondina and Old Nilpinna may have been caused by spring excavations and competition from the invasive mosquitofish *Gambusia holbrooki* Girard, 1859, respectively (Gotch et al., 2016). Extirpations of other fish species at Dalhousie have been attributed to the overgrowth of the native reed *Phragmites* as a result of decreased grazing pressure following the exclusion of livestock (Kodric-Brown et al., 2007). We also note that there are active springs at which only dead specimens of species have been collected, pointing to potential additional extirpations: for example, in certain spring groups within the Neales River and Francis Swamp complexes only empty shells of *Fonscochlea zeidleri* have been collected, with no evidence of live individuals (Ponder et al., 1989).

It is important to establish conservation priorities given the potential for further local, if not species-wide, extinctions in the SA GAB springs. Here, we ranked spring groups by the number of endemic species supported by each spring group and their degree of isolation. High-ranking locations mirror those proposed as conservation priorities in past studies, corroborating our findings. For example, McLaren *et al*. (1985) produced an inventory of fauna, flora, and funga to set out conservation priorities based on species diversity, the presence of rare species, “naturalness” (i.e., extent of interference by humans and cattle grazing pressure) and perceived vulnerability to degradation. Spring groups were then ranked by conservation priority. High-ranking spring groups in this assessment include those found at Dalhousie and in the Coward and Mt. Denison complexes of the Kati Thanda–Lake Eyre supergroup.

Further, Rossini *et al*. (2018) used a ranking system developed by Fensham and Price (2004) (modified here to produce endemicity rankings) to conclude that groups within the Dalhousie, Strangways, Francis Swamp, Billa Kalina, and Mt Denison complexes were of high conservation significance relative to other springs. The highest-ranking spring groups by endemicity in the current study presented here corresponded to all six of the above complexes, often down to the spring group level (Supplementary File S2). We stress that these trends do not suggest certain spring groups are more important than others or that insignificant springs exist, as has been implied in past environmental impact statements (Keane, 1997; Kinhill-Stearns, 1983). This is especially the case for locations that could be perceived as having low biological importance, such as the extinct Papu-ngaljuru / Primrose spring group, but have outstanding cultural significance to Aboriginal peoples due to associated Dreaming stories and its use as a major occupation site (Hercus and Sutton, 1985). Instead, our findings suggest that certain spring groups harbour high numbers of endemics relative to other locations (but remembering that insufficient sampling effort has occurred) and require targeted conservation efforts to preserve these short-range taxa.

Several conservation programs are currently in place which aim to directly improve the condition of SA GAB spring wetlands (Harris, 1992). Preliminary evidence suggests several of these practices have had a positive impact on SA GAB spring endemics. Springs are threatened by the presence of large-bodied and hard-hooved livestock which can graze on or trample wetland vegetation, foul the water through their faeces, and potentially severely reduce invertebrate biodiversity if cattle stocking levels are high; almost all SA GAB springs are subject to such pressure, particularly from cattle (Fensham et al., 2010; Gotch et al., 2016; Hutchinson and King, 1980; Kovac and Mackay, 2009). A combination of fencing and destocking has been successful in restoring endemic biodiversity of SA GAB springs in some cases, e.g., the endemic salt pipewort *Eriocaulon carsonii* subsp. *carsonii* F. Muell. in the Hermit Hill complex (Fatchen and Fatchen, 1993) and aquatic invertebrates at springs that were heavily damaged by stock (Kinhill-Stearns, 1984). The spring groups identified here as harbouring a large number of endemic species could represent future candidates for fencing and/or destocking initiatives.

As stated above, there is a large proportion of endemic taxa at several complexes within the SA GAB, namely those contained within the Dalhousie and Kati Thanda–Lake Eyre supergroups. The data presented here reiterates the need for targeted surveys in these locations to not only gather additional biodiversity data, but also to develop an understanding of population dynamics, habitat requirements, and accidental human-mediated translocations of fauna, flora, and funga per the National Recovery Plan (Fensham et al., 2010). The plan notably highlights the fact that the construction of a robust biodiversity inventory for GAB springs has been hindered by a lack of survey effort and taxonomic expertise. The utility of emerging environmental DNA techniques to capture a holistic picture of SA GAB spring biodiversity in a non-invasive manner would overcome several of these limitations (Beasley-Hall et al., 2023c; Saccò et al., 2022; Vörös, 2017; West et al., 2020); in this regard, the database developed as part of this study would assist in the selection of fieldwork locations for a pilot study in this regard. Once taxa are established as occurring within the SA GAB, monitoring efforts should be undertaken on a regular basis to develop deeper understandings of ecological knowledge as opposed to presence/absence information.

Although taxa reliant on GAB-fed springs are protected as a single ecological community under the EPBC Act, species-level listings are also far overdue. This is particularly the case for invertebrates dependent on the GAB. The majority of GAB molluscs and crustaceans meet the criteria to be listed as Critically Endangered (Rossini, 2020), but almost 90% of SA GAB invertebrates recorded here lack conservation listing at a global, federal, or state level, the bulk of which are either new species awaiting formal description or those completely lacking species-level identifications. This large knowledge gap is only exacerbated by a lack of understanding of the relationship between spring characteristics and biodiversity metrics (Fensham and Laffineur, 2022; Rossini et al., 2018) and indeed, relationships among spring ecological and hydrogeological characteristics themselves (Green and Berens, 2013; Love et al. (eds), 2013; Mudd, 2000). We hope that resources like the database developed here will inform the prioritisation of certain spring locations and taxonomic groups, ultimately leading to the listing of invertebrates and other spring endemics under relevant legislation on a per-species basis.

## 5 Conclusion

Here, we have presented the first review of biota supported by the Great Artesian Basin in South Australia. This dataset is an important resource we hope will facilitate future studies on GAB spring endemics, investigations into their population dynamics, basic biology, and taxonomy, and ultimately facilitate the listing of relevant taxa under state and federal environmental legislation. Such resources are essential given the multitude of threats currently facing springs and their biota, with potential extirpations of populations of at least 13 species in the SA GAB springs to date. We have also highlighted springs of particular conservation concern which may assist in determining future conservation priorities for this system. Our dataset has highlighted three major points about GAB spring biodiversity. Firstly, the majority of taxa reliant on these wetlands are invertebrates, and these animals are also the most poorly known and conserved in the GAB. Secondly, ecological communities reliant on artesian springs in SA are largely non-overlapping, irrespective of whether only endemic taxa, or endemics as well as “incidental” species, are concerned. The extinction or considerable disturbance of any spring group is therefore likely to lead to a considerable loss of biodiversity and/or genetic diversity. Finally, the artesian springs of SA have not been adequately sampled by past survey efforts and a considerable proportion of taxa likely remain to be documented, particularly in understudied locations such as the Munda / Lake Frome supergroup. There also remains a dearth of occurrence records for certain taxa for which taxonomic expertise has been lacking in the past, such as the wasps and allies (Greenslade, 1985). We recommend that datasets such as these are made publicly available with the capacity to be modified and updated on an ongoing basis, embodying the gold standard of digital asset storage.

## 6 Statements and Declarations

### 6.1 Data Availability Statement

All raw data associated with this paper are available via FigShare (Beasley-Hall et al., 2023a, 2023b).

## 6.4 Acknowledgements

The authors would like to acknowledge the Arabana Aboriginal Corporation, Travis Gotch, and The Friends of Mound Springs for their guidance and feedback.

